# Artificial-Cell-Type Aware Cell Type Classification in CITE-seq

**DOI:** 10.1101/2020.01.31.928010

**Authors:** Qiuyu Lian, Hongyi Xin, Jianzhu Ma, Liza Konnikova, Wei Chen, Jin Gu, Kong Chen

## Abstract

Cellular Indexing of Transcriptomes and Epitopes by sequencing (CITE-seq), couples the measurement of surface marker proteins with simultaneous sequencing of mRNA at single cell level, which brings accurate cell surface phenotyping to single cell transcriptomics. Unfortunately, multiplets in CITE-seq datasets create artificial cell types and complicates the automation of cell surface phenotyping. We propose CITE-sort, an artificial-cell-type aware surface marker clustering method for CITE-seq. CITE-sort is aware of and is robust to multiplet-induced artificial cell types. We benchmarked CITE-sort with real and simulated CITE-seq datasets and compared CITE-sort against canonical clustering methods. We show that CITE-sort produces the best clustering performance across the board. CITE-sort not only accurately identifies real biological cell types but also consistently and reliably separates multiplet-induced artificial-cell-type droplet clusters from real biological-cell-type droplet clusters. In addition, CITE-sort organizes its clustering process with a binary tree, which facilitates easy interpretation and verification of its clustering result and simplifies cell type annotation with domain knowledge in CITE-seq.

## 1 Introduction

Accurate cell type identification is critical to single cell analysis (Aevermann, et al., 2018; Klein and Treutlein, 2019). Surface markers are the most reliable biomarkers for cell type classification (Cui, et al., 2019; Spitzer and Nolan, 2016). Numerous studies in recent decades have used cell surface markers to isolate and characterize cellular populations with distinct biological functions and have discovered many novel cell types in the process (Ahmed, et al., 2019; Barcenilla, et al., 2019). Recent advancement in single cell RNA sequencing (scRNA-seq) technologies enabled direct cell identity assessment through dissecting single cell gene expression profiles. However, due to technical limitations, such as low efficiency in reverse transcription (Kharchenko, et al., 2014; Schwaber, et al., 2019), dropout events (sequencing fails to capture any transcript of active genes) (Li and Li, 2018; Macosko, et al., 2015) and low sampling rate (Macosko, et al., 2015; Stegle, et al., 2015), the RNA-derived cell identities may significantly deviate from the surface marker phenotyping result.

The recent introduction of cellular indexing of transcriptomes and epitopes by sequencing (CITE-seq) (Stoeckius, et al., 2017), which concurrently measures the abundance of both surface marker and messenger RNA (mRNA) in individual cells, enables simultaneous cell type identification based on surface marker and RNA profiles. In a CITE-seq dataset, surface markers are specifically labeled by antibody-derived tags (ADTs), which are individually barcoded. ADTs act as pseudo-transcripts and could be sequenced together with mRNA. CITE-seq has gained popularity in a wide range of biological and clinical studies, such as resolving tumor heterogeneity (Ortega, et al., 2017; Wagner, et al., 2019) and dissecting cell populations in complex tissues or organs (Cuevas-Diaz Duran, et al., 2017; Cui, et al., 2019; Klein and Treutlein, 2019; Landhuis, 2018; Schiller, et al., 2019).

**Figure 1A** and **1B** shows an example of cell type classification results of a peripheral blood mononuclear cells (PBMCs) CITE-seq dataset, using its RNA and surface marker (ADT) profiles separately. For RNA (**Figure 1A**), cells are clustered into separate groups using *scanpy* (Wolf, et al., 2018), based on the similarities in RNA profiles between cells; whereas for surface markers (**Figure 1B**), cells are clustered using Gaussian Mixture Model (GMM). We use manually annotated cell types, which are hand-curated with domain knowledge (Maecker, et al., 2012), as truth. In both plots, cells are embedded in 2-D planes, according to the tSNE transformation (Van Der Maaten, 2014) of their RNA or ADT profiles, respectively. Different cell types are highlighted with distinct colors. By comparing the classification results of both modalities, we observe that classification based on RNA profiles struggles to separate biologically similar but functionally unique cell types, such as cytotoxic T cells and helper T cells, or CD14^−^ monocytes and CD14^+^ monocytes, even though such cell types are distinctly separable in the surface marker space. A more detailed analysis between gene expressions and surface markers, such as the correlation analysis between them and RNA dropout analysis are provided in the Supplementary Figure 1. While GMM achieves a greater fidelity in the surface marker (ADT) space, it is still imperfect. In **Figure 1B**, GMM misclassifies a portion of the CD4^+^ T cells as some other cell types and fails to separate [iNK, mNK] and [C-mono, NC-mono] cell types into distinct clusters.

**Figure 1.**
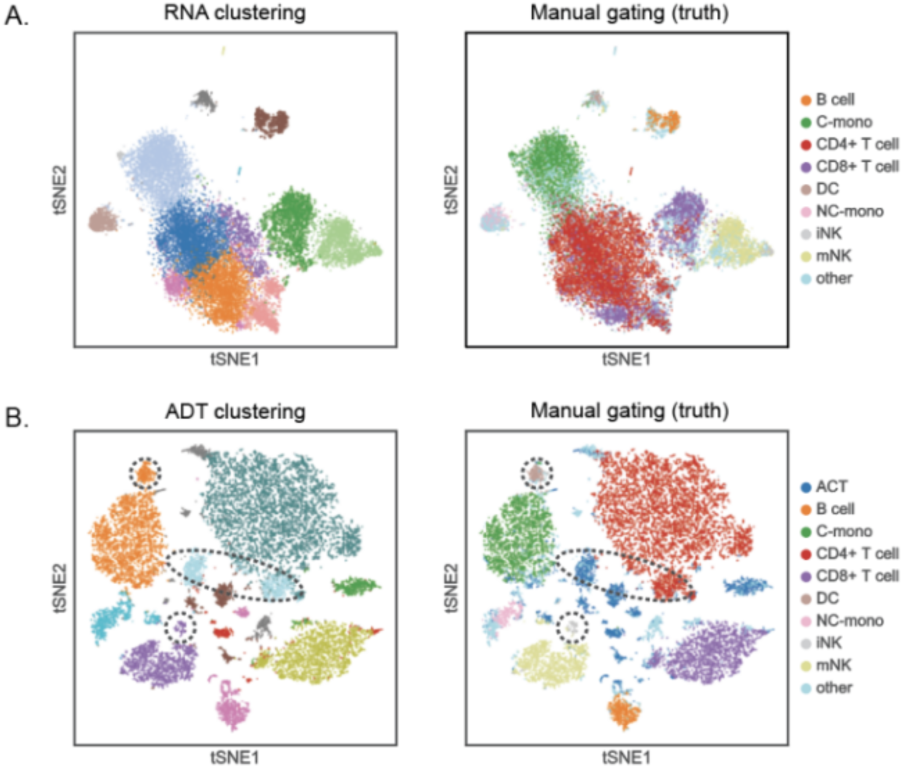
Clustering results of an example PBMC CITE-seq dataset. A) Droplets are clustered and arranged according to their RNA expression profiles. B) Droplets are clustered and arranged based on their surface marker (ADT) profiles. Ground truths in both plots are obtained through manual gating over the surface marker (ADT) space.

Automated cell type classification with surface markers in CITE-seq is further complicated by hard-to-remove multiplets. Multiplets, where more than one cell is found per droplet, are technical artifacts induced during library preparation (Macosko, et al., 2015; Stoeckius, et al., 2018). In single cell sequencing, ideally, a droplet is designed to capture just a single cell, which forms a singlet. In practice, it is impossible to avoid having two or more cells being encapsulated into a single droplet, which forms multiplets (see **Figure 2A**). Handling multiplets becomes an even more pressing problem when investigators super-load the library prep equipment, in order to cut reagent cost, eliminate inter-sample batch effect, or simply scale up samples in a single experiment. Multiplets confound cell type identification (McGinnis, et al., 2019; Wolock, et al., 2019). **Figures 2B** shows an example of multiplets creating artificial cell types in a PBMC CITE-seq dataset. When a multiplet contains cells of different types, it generates droplets with biologically impossible surface marker profiles. For instance, in the above example (**Figure 2B)**, there exists a sizable CD3^+^CD19^+^ droplet cluster which is rarely observed in PBMCs. Prior work has shown that such droplets are all multiplets and recommends subsequent removal of such droplets from downstream analysis (Xin, et al., 2019). In this work, we call cell types that truly exist as *bio*log*ical cell types* (BCT) and cell types created by CITE-seq multiplets as *artificial cell types* (ACT).

**Figure 2.**
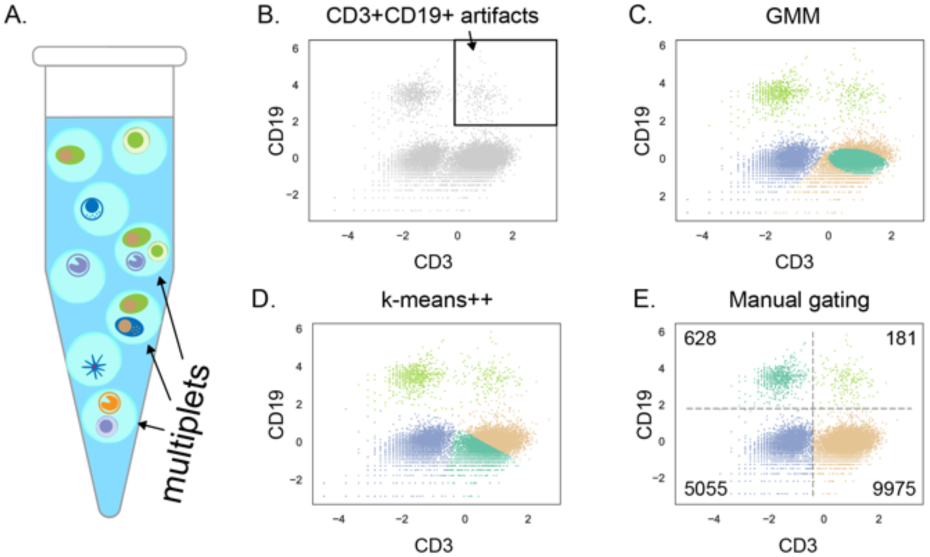
Demonstration of multiplets in CITE-seq and its impact on clustering methods. A) An example CITE-seq assay, which contains both singlets and multiplets. B) The example PBMC population displayed in the CD3:CD19 surface marker space. An ACT cluster in the pane is highlighted in the black box. C) and D), GMM and k-means++ clustering results with 4 clusters. E) Manual gating result, with the size of each cluster labeled in corners. The ACT cluster is much smaller in size than the three BCT clusters.

Since it is impossible to avoid the occurrence of ‘multiplets’ due to experimental limitations, we proceed to assess the impact of the multiplets on conventional clustering algorithms. **Figures 2C** and **2D** show the results of two popular clustering algorithms, Gaussian Mixture Model (GMM) and k-means++, for two surface markers, CD3 and CD19. With domain knowledge, the surface marker space should be divided into 4 quadrants with each quadrant containing a different cell type cluster, where top left are CD19^+^ cells, bottom left are CD3^−^CD19^−^ cells, bottom right are CD3^+^ cells and the top right are the joint CD3^+^-and-CD19^+^-cell ACT multiplets (**Figure 2E**). For this example, neither GMM nor k-means++ could isolate the top-right ACT cluster from the BCT clusters. This is due to the imbalance in cluster sizes between ACT and BCT clusters, where ACT clusters are significantly smaller than BCT clusters. The phenomenon where conventional clustering methods could fail when applied to datasets with mixing coefficient imbalances has been extensively studied and documented (Krawczyk, 2016; Lu, et al., 2019; Naim and Gildea, 2012; Xuan, et al., 2013).

Interpreting the clustering result is equally important (Kiselev, et al., 2019). Biologists have already established a mature pipeline to identify cell populations using a recursive selection process called gating, which is widely used in flow and mass cytometry analysis (Maecker, et al., 2012; Verschoor, et al., 2015). The gating process recursively selects a sequence of surface markers to stratify cells, until a homogeneous population emerges. The ordering of surface markers in gating is determined by domain knowledge. **Figures 3** illustrates an example gating strategy for PBMCs. For instance, to select the cytotoxic-T cells, which are marked by high expressions in both CD3 and CD8 and low expressions in both CD4 and CD19, a biologist would first select the CD3^+^CD19^−^ cell cluster in the CD3:CD19 scatter plot and then select the CD8^+^CD4^−^ cell cluster in the CD4:CD8 scatter plot (**Figure 3B**). Similar to how a gating strategy guides cell type annotation in manual gating, a good clustering framework should provide a comprehensive interpretation for its internal mechanics and facilitate biologists to annotate resulting cell clusters according to domain knowledge. Ideally, the interpretation should match the format of a gating strategy, such as the one demonstrated in **Figure 3A**.

**Figure 3.**
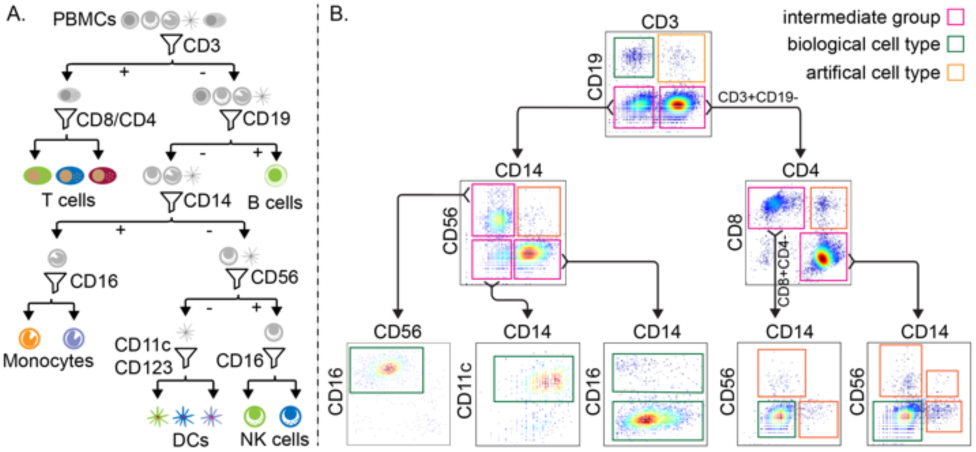
A) An example gating strategy for PBMCs. B) The gating process for the example PBMC dataset. Abbreviations: NK cells: natural killer cells; DCs: dendritic cells.

To address these challenges, we propose an interpretable, unsupervised single cell clustering algorithm, named CITE-sort, to systematically identify both BCTs and ACTs based on a recursive Gaussian Mixture Model framework. In each iteration, CITE-sort selects a low-dimensional surface marker subspace, generally defined by 1 or 2 surface markers, and performs clustering with GMM. In this manner, CITE-sort collapses numerous highly-imbalanced clusters in the original high-dimensional surface marker space into fewer yet more balanced hybrid clusters in lower-dimensional subspaces, which improves clustering performance. CITE-sort partitions the whole dataset into subpopulations through recursive bi-partitioning and constructs a binary sort tree for visualization, verification and guided cell type annotation. CITE-sort terminates when a subpopulation cannot be further divided, which creates a leaf node that is either an ACT or a BCT. The sort tree resembles a manual gating strategy and thus is comprehendible to biologists. The sort tree facilitates easy cell type annotations and accurate ACT cluster identifications. We applied CITE-sort to both real and simulated CITE-seq datasets and found that CITE-sort is able to achieve superior clustering performance compared to other popular clustering methods. Especially, CITE-sort is the only method that constantly and robustly segregate ACT clusters from BCT clusters. In addition, the sort tree generated by CITE-sort is consistent with existing gating strategies and provide rich biological insights.

## 2 Methods

### 2.1 Overlapping, imbalanced clusters in CITE-seq

CITE-seq uses a matrix to store the ADT counts of each surface marker in every droplet. While the ADT count of a surface marker does not quantify the absolute number of the marker proteins on a cell surface, it retains a strong correlation (Stoeckius, et al., 2017). Droplets that exhibit relatively large ADT counts of a specific surface marker, suggest that the cell(s) in these droplets have relatively high expression(s) of that surface marker. When multiple cells collide into a single droplet, forming a multiplet in the process, the final ADT counts of the droplet can be perceived as the sum of ADT counts over its member cells.

Similar to other surface marker quantification technologies, a CITE-seq dataset is pre-processed with the centered-log-ratio (CLR) normalization (Stoeckius, et al., 2017), prior to clustering. The formula of the CLR normalization is shown below:

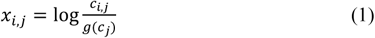

In **Equation 1**, *c*_*i,j*_ is the molecule count of marker (ADT) *j* in droplet *i* and *g*(*c*_*j*_) is the geometric mean of *c*_*j*_ across all droplets. All surface marker (ADT) scatter plots in this paper are CLR normalized.

CITE-seq datasets contain many imbalanced and overlapping clusters. ACT clusters are often in close proximity to multiple BCT clusters, similar to the example in **Figure 2B**. Compared to BCT clusters, ACT clusters have much smaller population sizes. In a CITE-seq experiment, the multiplet rate is often controlled at a moderate level for quality assurance. Previous work has shown that the proportion of multiplets increases as the number of cells in library prep increases (Macosko, et al., 2015; Stoeckius, et al., 2018; Xin, et al., 2019). Therefore, ACT droplets typically have non-negligible but still much smaller presences than BCT droplets, creating imbalance in cluster sizes.

In the CLR normalized surface marker space, an ACT cluster is always close to some BCT clusters. Each ACT droplet contains multiple cells of different types. Since the ADT counts of an ACT droplet equals to the sum of the ADT counts of its member cells, also because the logarithm dampens small-scale changes in the ADT counts (such as log(2 · *c*) = log(*c*) + log(2), with *c* ≫ 2), when individual cells merge into an ACT droplet, in the surface marker space, after CLR, the ACT droplet is positioned as slightly exceeding the highest coordinate in each surface marker dimension, across all of its member cells (Xin, et al., 2019). Mathematically, assuming ***C***_*D*_ is the set of cells included in an ACT droplet *D*, then the CLR-transformed surface marker count vector ***x***_*D*_ of *D* approximately equals

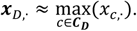

The produced ACT droplets, therefore, share similar ADT (CLR normalized) values to its member cells in many, but not all, surface marker dimensions. This creates the phenomenon where an ACT cluster is always close to some BCT clusters.

Aside from ACT clusters, BCT clusters may also share extensive similarities and form groups of closely positioned clusters in the surface marker space. Biologically, many cell types in a tissue share extensive common sections in their lineages. As a result, many of them share similar distributions in many surface markers. For instance, cytotoxic T cells (CD4^+^ T) and helper T cells (CD8^+^ T) are both T cells. As such, they have very similar, if not identical, ADT value distributions in many surface markers. Likewise, both T cells and B cells are lymphoid cells and their surface marker expression profiles also share significant similarities, albeit to a lesser extent than the helper and cytotoxic T cells (as T and B cells are more distant in the cell lineage tree).

Consequently, when observed in lower dimensions, especially when the most differentiating surface markers are excluded, BCT clusters often collapse with each other and merge into fewer clusters. **Figure 4** shows an example surface marker (ADT) distribution (CLR-transformed) of the previously-introduced PBMC dataset. In this dataset, when viewed in 1-D, many surface markers exhibit a mixture distribution of a few (mostly just two) Gaussian components. Also evident in **Figure 4**, these Gaussian components are seldomly well separated. In many surface marker dimensions, components are close to each other and share small overlaps.

**Figure 4.**
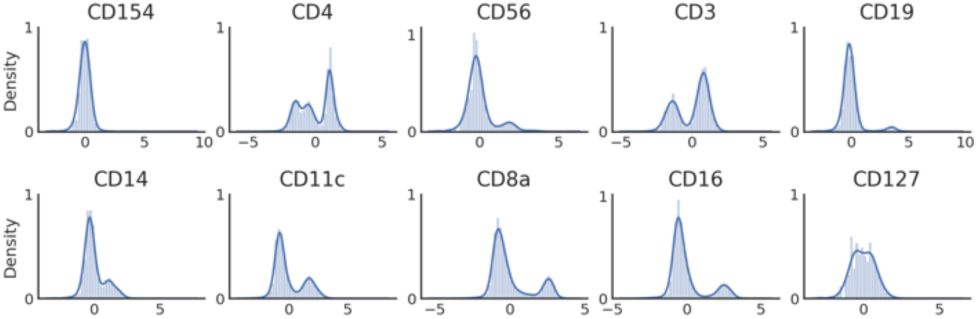
Distributions of CLR normalized ADT values of the example PBMC dataset on individual surface markers.

Combined with the property that ACT clusters are close to BCT clusters, in a CITE-seq dataset, there are many closely-positioned clusters in a very compact surface marker space. This increases the difficulty in separating them through clustering. Additionally, ACT clusters encompass much fewer cells than BCT clusters; and there could be many more ACT clusters than BCT clusters. According to Xin, et al. (Xin, et al., 2019), with a total of *k* BCT clusters, there could be as many as 2 ^*k*^ − *k* − 1 ACT clusters.

Altogether, a CITE-seq dataset is likely to share the following properties in the surface marker space: (1) it may contain a large number of clusters; (2) clusters vary dramatically in size; (3) clusters are not well separated and may contain similar distributions in individual dimensions.

### 2.2 Convergence of the EM algorithm on CITE-seq datasets

The expectation-maximization (EM) algorithm for GMM struggles to converge to the global optima in CITE-seq datasets. Inherently, in GMM, the EM algorithm does not consistently converge when (1) the dataset has high dimensions; (2) the cluster number is large; (3) clusters overlap and (4) there exists significant imbalance in the mixing coefficients. Let ***μ***\* denote the ground truth means of *K* Gaussian components in a *d*-dimensional Gaussian mixture dataset and let

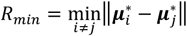

, where 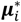 and 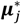 denote the ground truth means of component *i* and *j*, denote the minimum distance between the means of any two components. Zhao, et al., 2018 provides a convergence guarantee of the EM algorithm to the global optima, as long as the following condition is true:

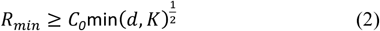

In **Equation 2**, *C*_*0*_ is a universal constant. **Equation 2** implies that it requires a greater distance between the means of components in a Gaussian mixture to provide the same convergence guarantee as the cardinality of dimensions and the number of clusters increase. Increasing dimensions or adding more components without increasing the distance between clusters will eventually undermine the convergence of the EM algorithm.

Naim and Gildea (Naim and Gildea, 2012) and Ma et al. (Ma, et al., 2000) have theoretically and empirically shown that having imbalanced mixing coefficients and overlaps between components also impedes the convergence of the EM algorithm. Specifically, Naim and Gildea (Naim and Gildea, 2012) showed that in the presence of overlaps between clusters, the condition number associated with EM increases (the convergence rate decreases) as the imbalance in mixing coefficients increases; hence a slower convergence of the EM algorithm.

CITE-seq datasets, unfortunately, often contain all four traits that hinders the convergence of the EM algorithm for GMM: there are many ACT clusters due to multiplets; ACT and BCT clusters often share overlaps; ACT clusters are much smaller in size than BCT clusters, which creates imbalance in mixing coefficients; and CITE-seq datasets usually have decent numbers of dimensions (often at the scale of ten or more). As a result, the EM algorithm exhibits poor performance at fitting a GMM model on a CITE-seq dataset, as shown in **Figure 2C** and **2D**.

### 2.3 Overview of CITE-sort

CITE-sort exploits a property of CITE-seq datasets where clusters of distinct cell types tend to collapsed into fewer Gaussian components in lower dimensions. Therefore, when inspected in a lower dimensional subspace, the distribution can be fitted with much fewer components. Working with fewer components in a low dimensional space improves the likelihood of the EM algorithm reaching to the global optima (recall **Equation 2**). For example, when viewed in 1-D, the four Gaussian components in **Figure 5C** collapses into only two components, as **Figure 5A** shows. When directly applying the EM algorithm on all four components in the 2-D space, the EM algorithm fails to converge to the global optima and is unable to separate the four clusters, as with the example in **Figure 2C**. However, by applying one-dimensional GMM iteratively on CD3 (**Figure 5A**) and then on CD19 (**Figure 5B** and **5C**) in a two-step fashion, the EM algorithm is able to reliably crop out all four clusters, as displayed in **Figure 5**. When clustering on CD3, the two Gaussian components are more balanced in size; hence leading to a more consistent convergence for the EM algorithm. When processing the CD3^+^ component on CD19 (**Figure 5C**), although there still exist non-trivial mixing coefficient imbalance between the CD3^+^CD19^+^ component (an ACT cluster) and the CD3^+^CD19^−^ component (a BCT cluster), compared to the original two-component, 4-cluster scenario in **Figure 5A**, which harbors mixing coefficient imbalance between the CD3^+^CD19^−^ ACT cluster and all three other BCT clusters, the imbalance is now restricted between just a single pair of components. Combined with having fewer clusters in the mixture and operating in lower dimensions, the EM algorithm is able to converge accurately and separate the CD3^+^CD19^+^ and the CD3^+^CD19^−^ cluster.

**Figure 5.**
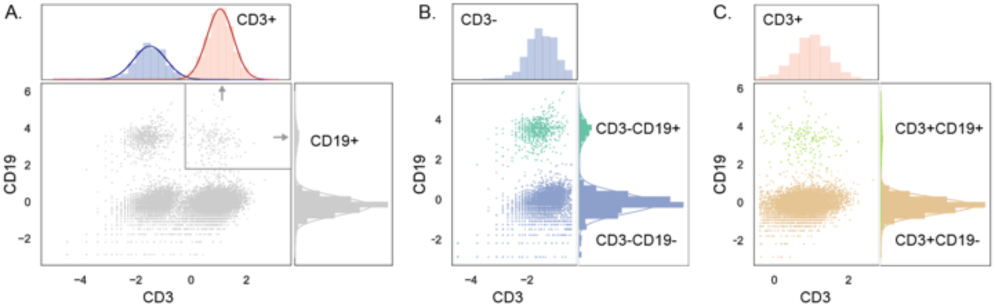
Demonstration of CITE-sort fitting GMM models in lower dimensions. A) Fitting a GMM model in single dimensions. B) and C) Fit GMM models with CD19 on CD3^-^ and CD3^+^ subpopulations from the previous step, respectively.

Overall, CITE-sort clusters droplets by iteratively fitting GMM models in low dimensions. In each iteration, CITE-sort selects the lowest dimensional subspace where droplets can still be distributed into separate hybrid clusters. If there exist multiple qualifying surface marker subspaces with equal cardinalities, CITE-sort prioritizes the surface marker subspace whose GMM model returns the largest likelihood in describing the data distribution. CITE-sort repeats the low-dimensional clustering process on each child cluster, until no cluster can be further subdivided. In this manner, CITE-sort reduces the number of clusters, the cardinality of the dimension, and the degree of imbalance among clusters in each clustering step and maintains high quality clustering throughout the entire process.

For simplicity, CITE-sort uses binary partition: in each iteration, CITE-sort picks the largest hybrid cluster as one part and merges all other components together as the other part. CITE-sort then repeats the low-dimensional clustering process on each part, until it can no longer be further divided. We call the binary tree emerged from the iterative bi-partitioning process *the sort tree* (sort for cell sorting).

### 2.4 Clustering in low-dimensions

When clustering in low dimensions, clusters may not always perfectly stack on each other and form perfect Gaussian components as the example in **Figure 6A.** Sometimes, the density distribution of a surface marker cannot be decomposed into two perfectly disjoint Gaussian components. Rather, it requires multiple components, some of which overlap with each other. CITE-sort merges Gaussian components that share significant overlaps, such as the example distribution shown in **Figure 6A**. CITE-sort computes the Bhattacharyya distance (Bouveyron, et al., 2019; Hennig, 2010) to measure the degree of overlap between two components. Given a surface marker subspace *S* and two Gaussian components *a* and *b* in *S*, Let *m*_*a*_, *m*_*b*_ denote the surface marker mean vectors of *a* and *b*, and Σ_*a*_, Σ_*b*_ denote the corresponding covariance matrices of a and b. The Bhattacharyya distance (Bouveyron, et al., 2019; Hennig, 2010) between *a* and *b* equals:

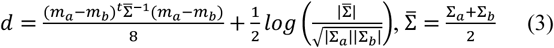

**Figure 6.**
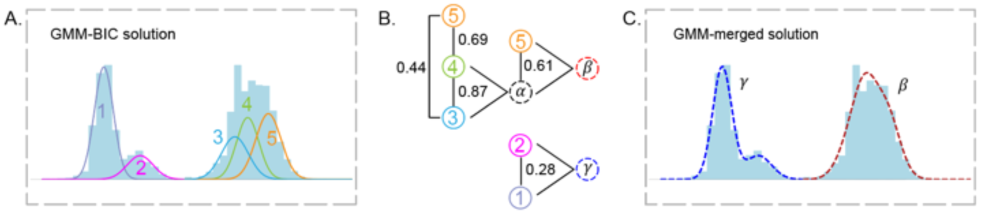
An example of merging Gaussian mixture components. A) Illustration of the 5 original Gaussian components in a 1-D GMM model. B) Merging components into component complexes. C) The final component complexes after merging. The dashed lines represent the density curves of the two merged component complexes *β* and *γ*.

If he Bhattacharyya distance between a pair of components falls below a parametrized threshold *ε*, then the two components are merged into a component complex. After merging, the same process is repeated on all pairs of component/component complexes, until no pairs of component/component complex can be merged. **Figure 6B** demonstrates the hierarchical merging process of combining Gaussian components of **Figure 6A**. There are 5 Gaussian components. Component pairs, [1, 2] and [3, 4], are first merged into a component complex *α* and *β*, respectively. Then *α* is merged with component 5, becoming *γ. β* and *γ* do not share sufficient overlap. Hence, they remain two separate component complexes. **Figure 6C** shows the final component complexes, *β* and *γ*, in the density histogram plot. CITE-sort sets *ε* to 0.1 by default.

### 2.5 Constructing the sort tree

As CITE-sort iteratively subdivide droplets into smaller subpopulations, CITE-sort records the surface marker(s) involved in each division in the sort tree. Each node in the sort tree represent a subpopulation. The root node represents the whole dataset. The label of each node denotes the surface markers employed in further subdividing the subpopulation. CITE-sort stops subdividing a node if 1) the population size falls below a parametrized minimum cluster threshold *θ*, or 2) the subpopulation cannot be subdivided into more than one component/component complex in all sub-spaces.

While CITE-sort prefers clustering in low dimensions, not all surface marker subspaces are qualified for subdivisions. Many surface subspaces have a single component complex left after merging. Without more than one independent component/component complex, the subdivision process comes to a halt. To avoid terminating the iterative clustering process prematurely, CITE-sort skips surface marker subspaces with a single component complex (or if there are no multiple components to begin with). When there exist multiple qualifying surface marker subspaces with equal cardinalities, CITE-seq prioritizes the surface marker subspace whose GMM produces the highest score in goodness of fit measurement. CITE-sort computes the likelihood of the fitted GMM in describing the distribution of the data and selects the surface marker subspace with the largest likelihood for further division.

The sort tree facilitates accurate annotations of cell types, justifiable verification of the clustering result, and accessible interpretation of the model. The sort tree displays three items at each node: the surface marker(s) chosen for clustering and bi-partition; the distribution of the current droplet subpopulation in the chosen surface marker subspace; and all component complexes derived from the subpopulation. The two subpopulations after bi-partition are highlighted separately with matching-color edges in the sort tree to emphasize their sorting trajectory. An example sort tree is provided in **Figure 9A** in the Result section.

**Figure 7.**
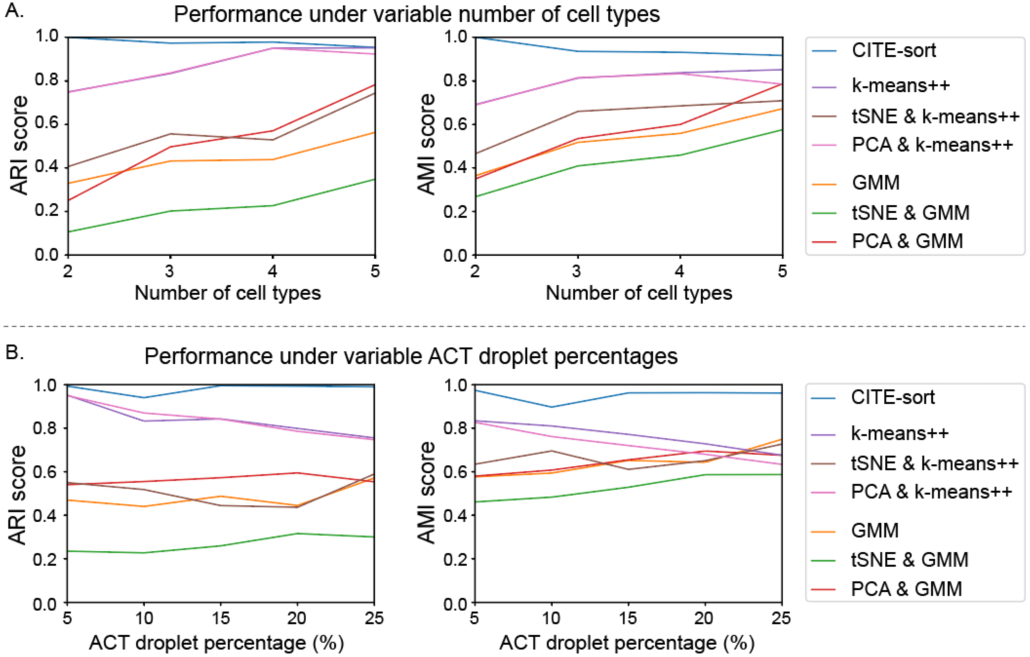
Performances of CITE-sort compared against 6 other common clustering methods under A) variable number of cell types (fixing ACT droplet percentage at 5%) and B) variable ACT droplet percentages (fixing the number of biological cell types at 4).

**Figure 8.**
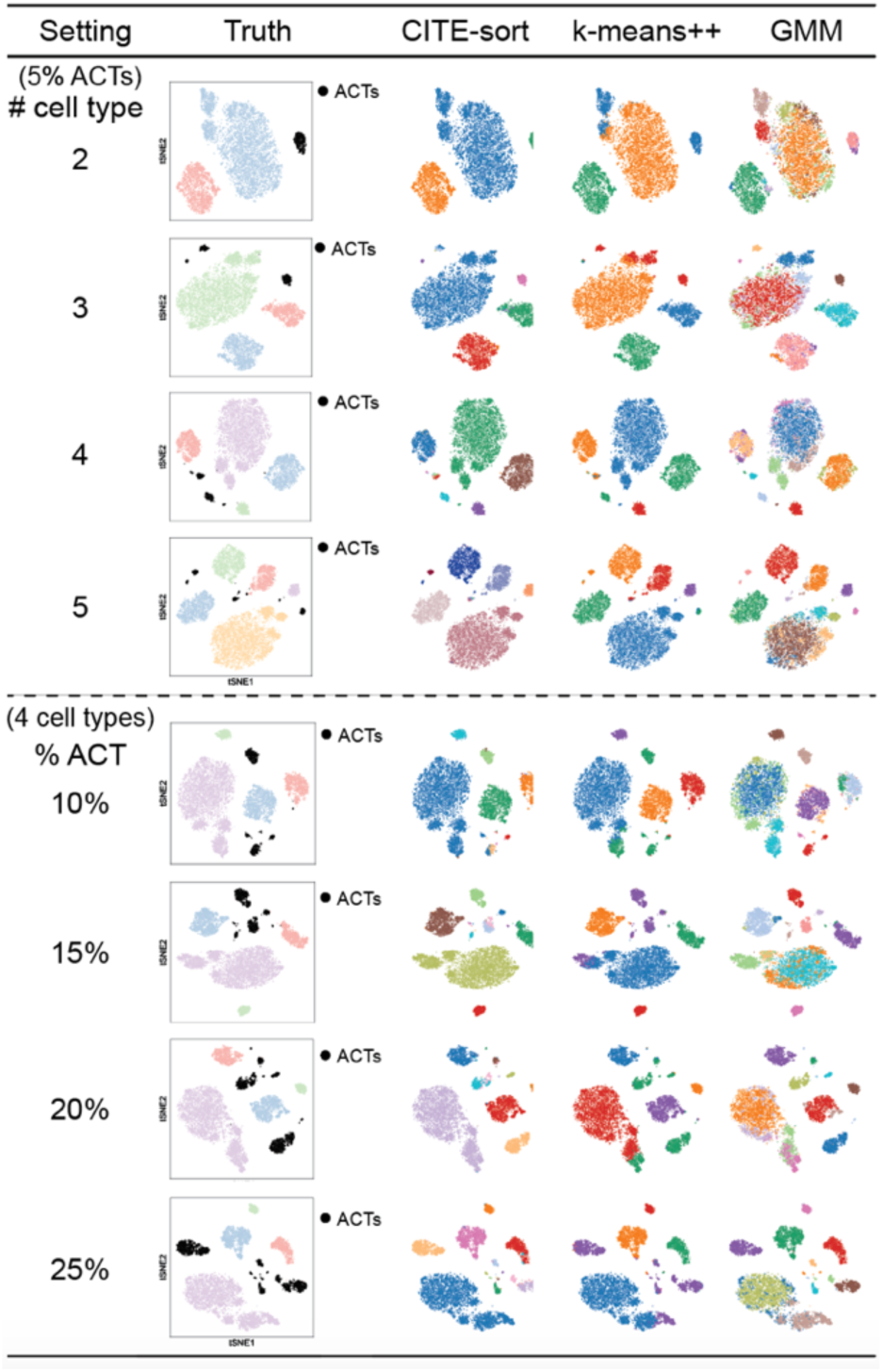
Clustering results of CITE-sort, k-means++ and GMM with regard to 8 simulation datasets. Different colors denote distinct clusters.

**Figure 9.**
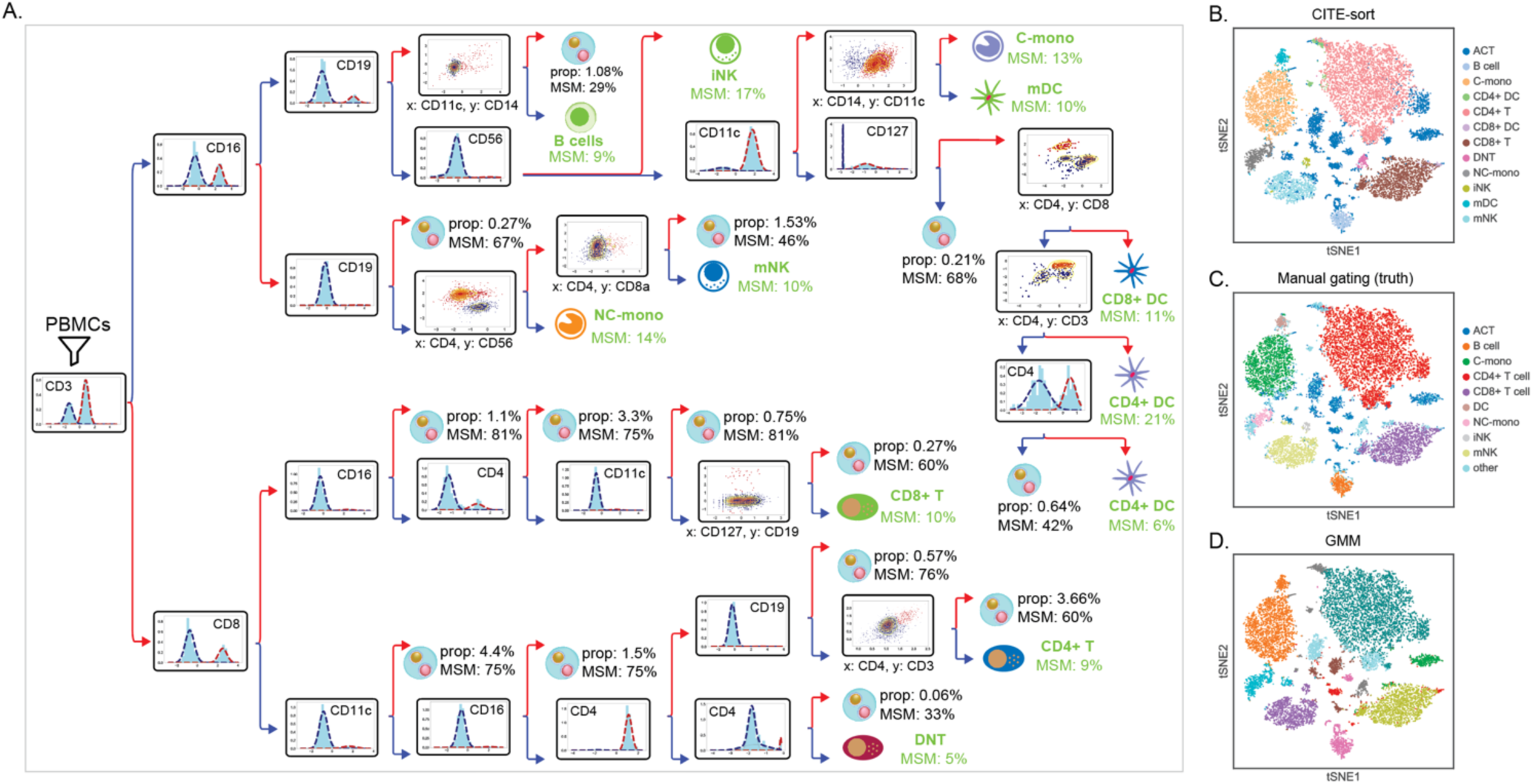
A) The single cell sort tree of PBMC_16K. Each node represents a subpopulation. The title of each inner node represents the surface markers subspace for subdivision. Red and blue colors represent the two parts in bi-partition. Edges are colored accordingly. Leaf nodes are hand-curated and are annotated with domain knowledge. Cell types that should not exist are labeled as suspect ACT clusters. Suspect ACT clusters are characterized by their population percentages in the overall dataset (denoted by ‘prop’) and their MSM percentages (denoted by ‘MSM’). Abbreviations: iNK: intermediate NK cells; mNK: vast majority of NK cells; C-mono: classical monocytes; NC-mono: non-classical monocytes; mDC: myeloid DC; DNT: double negative T cells. B) CITE-sort clustering result. C) Manual clustering result. D) GMM clustering result. All clustering results are projected over the tSNE-transformed 2-D surface marker pane.

The sort tree lets end users to inspect the distribution in the surface marker subspace at each stage and assess the fidelity of the clustering decisions made by CITE-sort. The sort tree also enables users to follow the sorting trajectory of any terminal cell population and register surface markers where the population have high or low levels of expressions. End users can then compare the sorting trajectory of each terminal cell population with existing gating strategies and annotate cell types according to domain knowledge.

### 2.6 Pseudo code

Let ***D*** = {***x***_l_, …, ***x***_*N*_} denote a CLR-transformed ADT dataset. Let *N* denote the number of droplets and ℱ = {*f*_1_, …, *f*_*M*_} denote the *M*-dimensional surface marker space of the dataset. CITE-sort takes ***D*** as input and returns the root node of the binary sort tree 𝒯. Each node in 𝒯 stores the surface marker subspace selected for clustering, the current subpopulation of the node and pointers to children nodes (leaf nodes point to NULL). The pseudo code of CITE-sort is provided in **Algorithm 1**.

#### Algorithm 1. The pseudo code of CITE-sort.

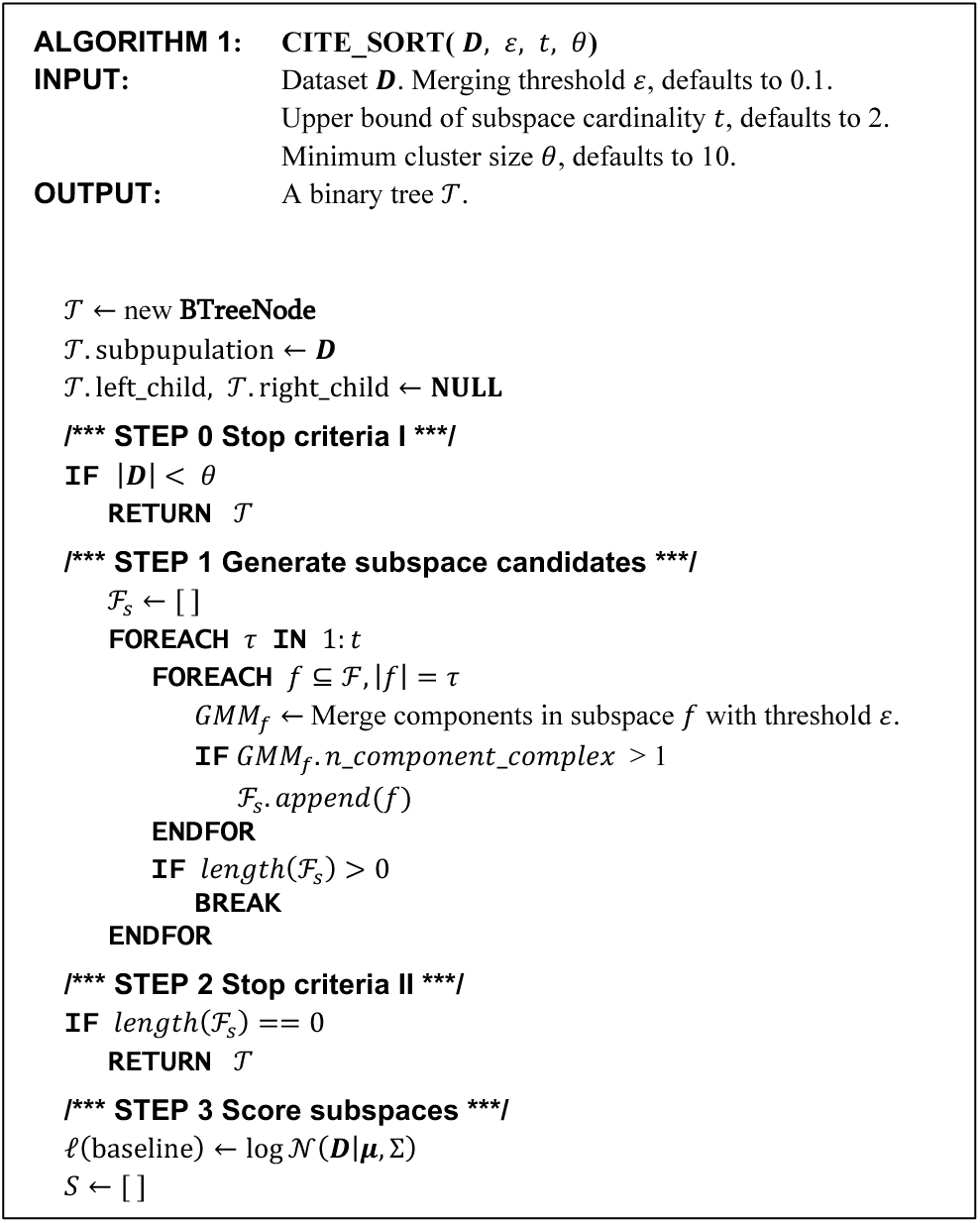
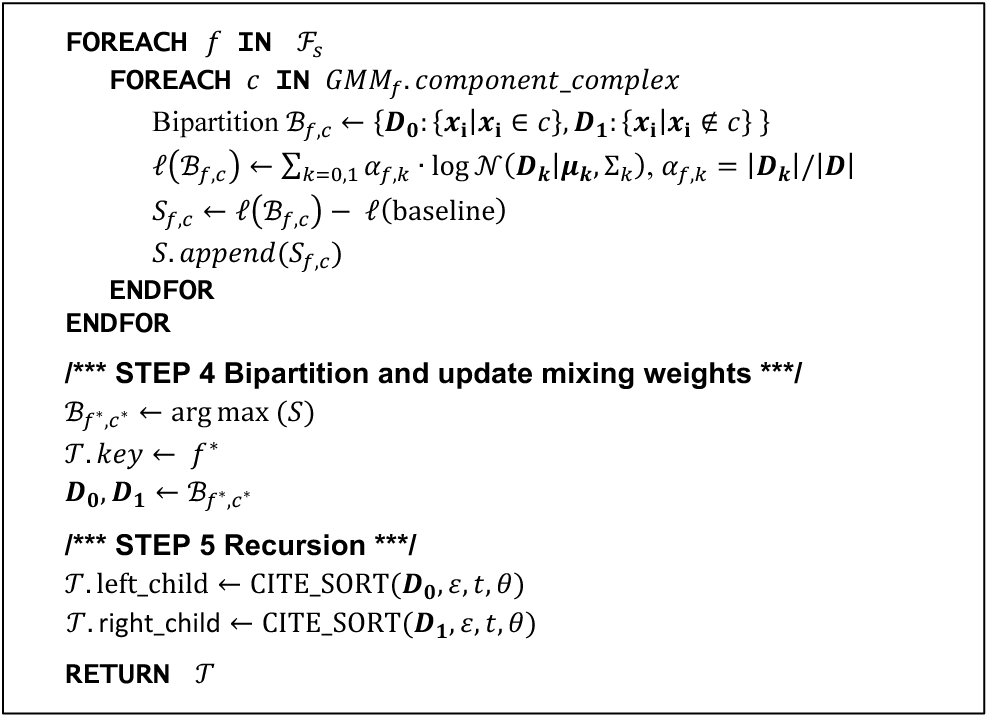

## 3 Datasets

### 3.1 Real Datasets

We tested CITE-sort with three in-house PBMC CITE-seq dataset as well as 5 public CITE-seq datasets. For the three in house datasets, cells were drawn from two healthy donor described in a previous study (Sun, et al., 2019). The cell assays were prepared using the 10x Genomics platform with Gel Bead Kit V2, and was subsequently sequenced on an Illumina HiSeq with a depth of 50K reads per cell. We measured ten surface markers on every cell. They were: CD3, CD4, CD8, CD11c, CD14, CD16, CD19, CD56, CD127 and CD154. We performed downstream analysis with CellRanger-3.0. Public datasets include three PBMC datasets, one marginal zone B-cell tumor (MALT: mucosa-associated lymphoid tissue) dataset and one cord blood mononuclear cells (CBMC) dataset (Stoeckius, et al., 2017). Detailed information of all real datasets is summarized in **Table 1**.

**Table 1.** Overview of CITE-seq datasets.

| Dataset | # Droplets | # ADTs | Source |
| --- | --- | --- | --- |
| PBMC_1k | 713 | 17 | 10X Genomics |
| PBMC_1k_b | 1388 | 10 | In house |
| PBMC_2k | 1966 | 10 | In house |
| PBMC_5k | 5247 | 32 | 10X Genomics |
| PBMC_8k | 7865 | 17 | 10X Genomics |
| MALT_8k | 8412 | 17 | 10X Genomics |
| CBMC_8k | 8617 | 13 | GSE100866 |
| PBMC_16k (with cell hashing) | 15839 | 10 | In house |

10X Genomics datasets are available at https://support.10xgenomics.com/single-cell-gene-expression/datasets.

We focus our evaluation of CITE-sort with the PBMC_16k dataset. For the PBMC_16k dataset, in addition to CITE-seq, we also performed cell hashing (Stoeckius, et al., 2018) with four hashtags. Cell hashing enables technicians to increase the population of cells in a single scRNA-seq run, and it facilitates analytical identification of a significant fraction of multiplets. Previous work has demonstrated analytical classification of putative homogeneous droplet populations into ATCs and BTCs with cell hashing (Xin, et al., 2019). In a cell hashed scRNA-seq dataset, droplets are categorized into either multi-sample multiplets (MSM) or single-sample droplets (SSDs). Prior work (Xin, et al., 2019) has established that ACT clusters have MSM percentages approaching and exceeding 1 − 1/*H*, where *H* is the number of hashtags used in cell hashing (assuming even cell distributions among samples in cell hashing). For the PBMC_16k dataset, we used *H* = 4 hashtags. If a putative homogeneous droplet cluster in PBMC_16k has a MSM ratio approaching or exceeding 75%, then it is highly likely to be an ACT cluster. We used GMM-Demux (Xin, et al., 2019) to classify droplets into MSMs and SSDs.

It is important to establish the evaluation of CITE-sort with cell hashed CITE-seq datasets. The MSM ratio of a cluster serves as an important signature of whether the cluster is an ACT or BCT. The ratios of MSMs in suspicious ACT clusters can be used to validate the efficacy of CITE-sort in separating ACT clusters from BCT clusters: high MSM content in suspicious ACT clusters and low MSM content in BCT clusters suggest that CITE-sort achieves high precision in isolating ACT clusters from BCT clusters; while the opposite indicates that the suspect ACT clusters have absorbed excessive BCT droplets because of poor clustering performance. Given that PBMC_16k is the only available cell-hashed CITE-seq dataset, we focus the evaluation of CITE-sort over the PBMC_16k dataset.

### 3.2 Simulation Datasets

We generated a number of CITE-seq simulation datasets according to the droplet formation model described in Xin, et al. (Xin, et al., 2019). Specifically, we assumed that cells were randomly distributed into a finite number of droplets. We prepared synthetic cells by sampling droplets from the in-house PBMC dataset. All synthetic cells are hand-curated into one of the 5 biological cell types through manual gating, which is summarized in Supplementary Table 1. Synthetic cells are randomly distributed to droplets. If a droplet is assigned with a single cell, then the raw ADT (yet to be CLR normalized) profile of the droplet equals to the raw ADT counts of its member cell. Otherwise, if a droplet is assigned with multiple cells, the raw ADT profile of the droplet equals to the sum of raw ADT counts over all member cells. Empty droplets are removed after simulation.

Droplets are labeled according to their member cell(s). ACT droplets that have different cell type compositions are assigned different labels. With *M* biological cell types in a simulation, there could be as many as 2^*M*^ − 1 distinct droplet labels, including M BCT labels and up to 2^*M*^ − M − 1 ACT labels. The droplet labels from simulation are used as the ground truth.

We performed two sets of simulations. In the first set, we controlled the percentage of ACT droplets at 5% and gradually increased the number of biological cell types in each simulation, from 2 to 5. In the second set, we fixed the number of biological cell types at 4 but gradually increased the ACT droplet percentage, from 5% to 25% with a 5% increase per step.

## 4 Results

### 4.1 Accuracy evaluation on simulated datasets

We compared CITE-sort against 6 other clustering methods: k-means++, k-means++ with PCA, k-means++ with tSNE, GMM, GMM with PCA, and GMM with tSNE. For both PCA and tSNE based clustering, we reduced the dimensions of each dataset to three. For k-means++, we used the elbow method with inertia (Bholowalia and Kumar, 2014) to select the number of clusters *K*; for GMM, we used the Dirichlet Process to infer *K* from the data (Blei and Jordan, 2006; Görür and Rasmussen, 2010).

We benchmark CITE-sort against the other six clustering methods on both sets of simulation datasets. We compare the clustering outcome of each clustering method based on both the Adjusted Rand Index (ARI) score and the Adjusted Mutual Information (AMI) score against the ground truth. As shown in **Figure 7**, CITE-sort has the best performance across the board in both experimental settings. Especially, when measured in AMI score, CITE-sort always leads other methods by at least 0.1 on average. Previous work (Romano, et al., 2016) has shown that AMI creates more accurate measurements than ARI, under scenarios where the cluster sizes are highly unequal in ground truth. Therefore, maintaining a commanding lead in AMI scores under low ACT droplet percentages proves that CITE-sort outperforms other clustering methods when clusters contain significant mixing coefficient imbalances. k-means++, GMM and their variants, on the other hand, have much inferior AMI scores under low ACT droplet percentages.

The clustering results of each simulation dataset is presented in **Figure 8**. Comparing to the other 6 clustering methods, CITE-sort produces clustering results that are most consistent with the ground truth. The reason that the clustering accuracy of GMM and K-means++ increases when more cell types are added to the simulation under a fixed ACT percentage threshold (5%) is that ACT droplets are distributed into a rapidly increasing number of ACT clusters (the number of ACT clusters grows exponentially as more cell types are added). With much more diluted ACT clusters, the influence of ACT clusters in disrupting the separation of BCT clusters decreases. Hence GMM and K-means++ register better performances. Eventually, as ACT clusters become increasingly sparse, we estimate that the performances of GMM and K-means++ will climb to the same level as CITE-sort. Nevertheless, to reduce the ACT percentage, technicians will have to drastically limit the number of cells loaded in a cell assay, which significantly limits the applicability of GMM and K-means++ in real datasets.

### 4.2 Constructing sort trees in real datasets

**Figure 9A** demonstrates the sort tree of the PBMC_16k dataset. In this dataset, clustering is performed in either 1-D or 2-D surface marker subspaces in each step. Depending on the dimension of the surface marker subspace, each inner node is visualized with either a histogram or a 2-D scatter plot. In each plot, we use red and blue to highlight the two parts after bi-partition and assigned colors to edges in the sort tree correspondingly. Each leaf node is annotated according to a standard PBMC gating strategy (Maecker, et al., 2012). We labeled leaf nodes that cannot be explained by the standard gating strategy as suspect ACT clusters. Leaf nodes are supplemented with its population percentages and MSM percentages. As mentioned in Section 3.1, according to GMM-Demux, droplet clusters with MSM percentages approaching and exceeding 75% (*H* = 4 in this dataset) are ACT droplet clusters. We observe that most suspect ACT clusters have high MSM percentages. Overall, we find an estimated 19.3% of all droplets that are ACT droplets.

The sort tree of PBMC_16k is highly consistent with existing gating strategies (see **Figure 3**). Compared to manual gating (**Figure 3**), CITE-sort achieves greater clustering resolutions. In particular, CITE-sort subdivides NK cells into majority NK cell and intermediate NK cell subtypes; monocytes into classical-monocytes and nonclassical-monocytes; and DCs into CD4^+^ DCs, CD8^+^ DCs and myeloid DCs. CITE-sort also splits double negative (CD4^−^CD8^−^) T cells from other T cells. Such resolution is hard to guarantee in manual gating. The resemblance between the sort tree and common gating strategies validates the clustering logic of CITE-sort. GMM and k-means++, on the other hand, lacks interpretability and thus renders their clustering results more difficult to verify.

**Figure 9B** presents the CITE-sort clustering result of PBMC_16k in a 2-D tSNE plot. Compared to GMM (**Figure 9C**), CITE-sort clustering is more consistent with human-supervised manual clustering (**Figure 9D**). Further comparison shows that major cell types in PBMC have highly consistent scopes in both CITE-sort (**Figure 9B**) and manual gating (**Figure 9D)** clustering results, showcasing the accuracy of CITE-sort in cell sorting.

**Figure 10** illustrates the effectiveness of CITE-sort in segregating ACT clusters. In **Figure 10B**, we combine all hand-curated ACT clusters into a single ACT class. We use GMM-Demux to classify droplets into MSMs and SSDs (**Figure 10A)**. Comparing **Figure 10A** and **10B**, we observe that the combined ACT super cluster accurately outlines major MSM rich regions. As previously discussed, high MSM concentrations in suspect ACT clusters and low concentration of MSMs in BCT clusters proves the accuracy of CITE-sort in segregating ACT clusters from BCT clusters.

We also compared the ACT annotation over CITE-sort against an RNA-based doublet identification method, scrublet (Wolock, et al., 2019). The classification result of scrublet is shown in **Figure 9C**. Scrublet identifies doublets by analyzing the RNA expression profile of a scRNA-seq experiment. It simulates artificial doublets from the dataset and checks if a droplet has an expression profile that matches the expression profile of any simulated doublet. Comparing to **9B, Figure 9C** only highlights a fraction of the multiplets (19.3% vs 6.4%) and completely misses a few doublet clusters (highlighted in circles). Since it leaves a large number of ACTs unidentified, it denies the possibility of clustering droplets after removing all ACTs. Removing all MSMs is also insufficient. Not all droplets in ACT clusters are MSMs while not all droplets in BCT clusters are SSDs. Removing MSMs does not completely erase ACT clusters. Other doublet removal tools, such as GMM-Demux requires pre-preparation of droplets into clusters and can only checks if an entire cluster is an ACT cluster. Overall, CITE-sort is the only ACT-aware CITE-seq clustering method.

**Figure 10.**
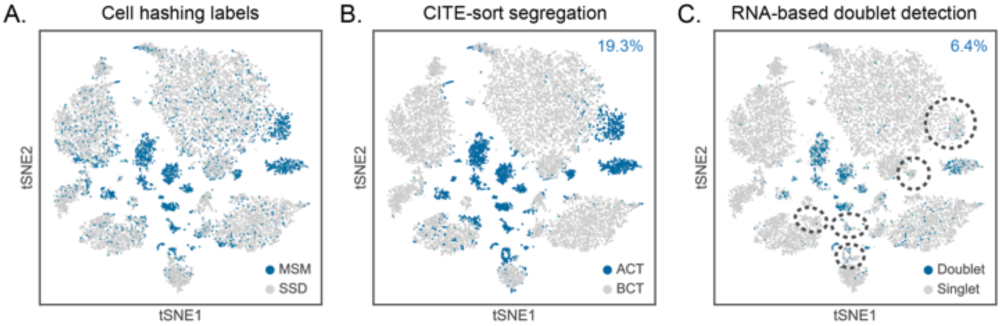
tSNE visualization of A) Suspect ACT clusters identified by CITE-sort, B) MSMs identified through cell hashing, and C) doublets identified by scrublet. The x and y axis are the first and second dimensions respectively given by tSNE. Percentages of ACTs/doublets identified by CITE-sort/scrublet are presented in the top right corner.

In addition to PBMC_16k, we also tested CITE-sort on the other 7 CITE-seq datasets. We benchmarked CITE-sort against GMM and its variants. Detailed description of each method is provided in the Supplementary Method 1. Unfortunately, without cell hashing, we are unable to benchmark the accuracy of ACT classifications by CITE-sort. Instead, we measure the likelihood of the final model in describing the distributions of the datasets. The likelihood comparison result is presented in **Figure 11A**. As shown in the figure, CITE-sort achieves the highest likelihood in all datasets. **Figure 11B** illustrates the execution time comparison between CITE-sort and GMM methods. Overall, CITE-sort terminates in comparable time against other methods.

**Figure 11.**
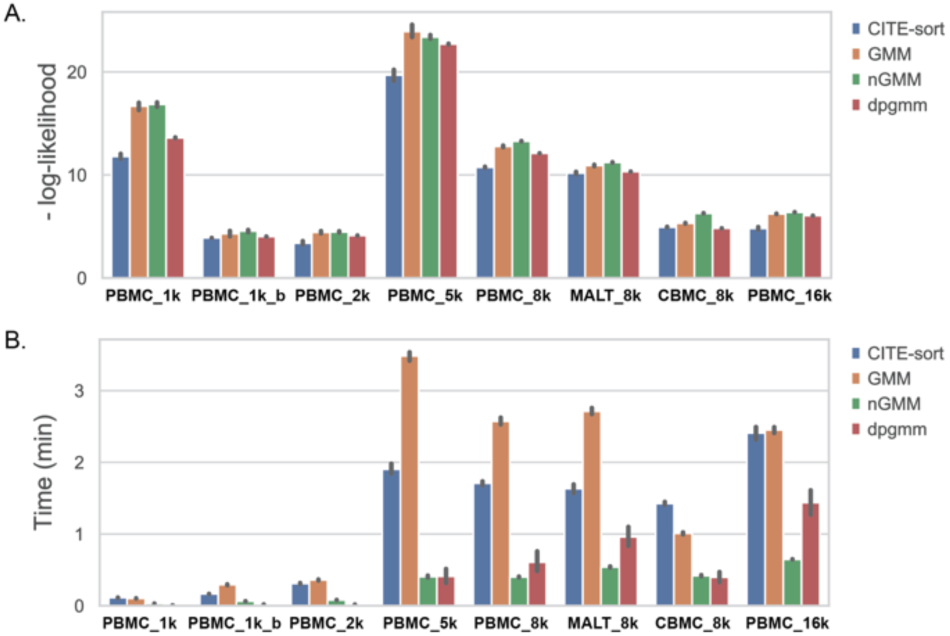
Performance comparisons of CITE-sort against GMM and its two other variants. Benchmarks includes A) negative log-likelihood and B) execution time comparison

## 5 Discussion

CITE-sort is the first ACT-aware surface marker clustering method for CITE-seq datasets. CITE-sort specifically targets datasets where clusters in high-dimensional space collapse into fewer, more balanced clusters in low dimensional subspaces. CITE-sort is not a universal clustering method. While it is possible to construct datasets where CITE-sort produces inferior clustering results than GMM; or CITE-sort has to scan through a large number of low dimensional subspaces before finding a qualifying candidate, in real CITE-seq datasets, most cell types collapse into very few Gaussian components when viewed in any single or dual-surface-marker subspaces. In fact, popular manual gating tools, such as FlowJo (Tree Star, 2020) and Kaluza (Beckman Coulter, 2020) are built upon this observation where cells are always grouped into few Gaussian clusters when viewed in low dimensions (usually in 2-D). Given these characteristics of CITE-seq datasets, CITE-sort is able to scale to higher dimensional surface marker spaces, even as the number of surface markers grows up to 100 and beyond in future CITE-seq datasets.

## 6 Conclusion

In this paper, we introduced CITE-sort, an interpretable clustering framework for CITE-seq datasets, which groups droplets into clusters in an unsupervised manner. CITE-sort is robust to artificial cell types that stem from multiplets. CITE-sort is the first clustering method for CITE-seq that is aware of artificial cell types. CITE-sort generates biologically meaningful interpretations to its clustering results. We applied CITE-seq on both real and simulated CITE-seq datasets and show that CITE-sort not only outperforms canonical clustering methods in accuracy, but also generates a single cell sort tree, which helps in annotating cell types, validating clustering results and identifying artificial-type droplet clusters.

