## Supplementary Figure 1, Supplementary Table 1, Supplementary Method 1 for "Artificial-Cell-Type Aware Cell Type Classification in CITE-seq"

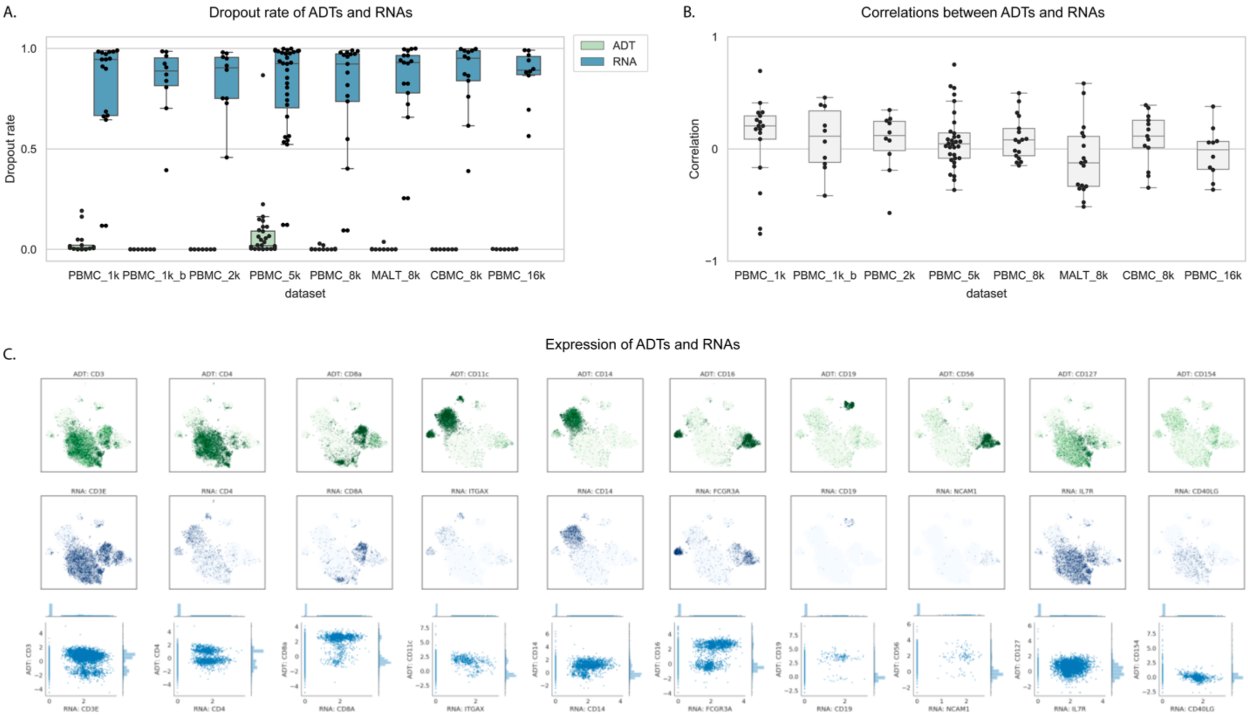


**Supplementary Figure 1**

**Analysis between RNAs and surface markers in CITE-seq.**

A) Dropout rates of RNAs and surface markers (ADT) in eight CITE-seq datasets. B) Pearson correlation between normalized RNA expression and CLR-transformed ADT counts in eight CITE-seq datasets. We normalize RNA counts to 10,000 reads per cell and log-transform the result. C) CLR-transformed ADT counts (upper) and normalized RNA expression (middle) are displayed in 2-d embedding space. The x and y axis are the first and second dimensions respectively given by tSNE based on normalized RNA expression profiles. The bottom panel shows the co-expression patterns between ADTs and RNAs in PBMC_16k.

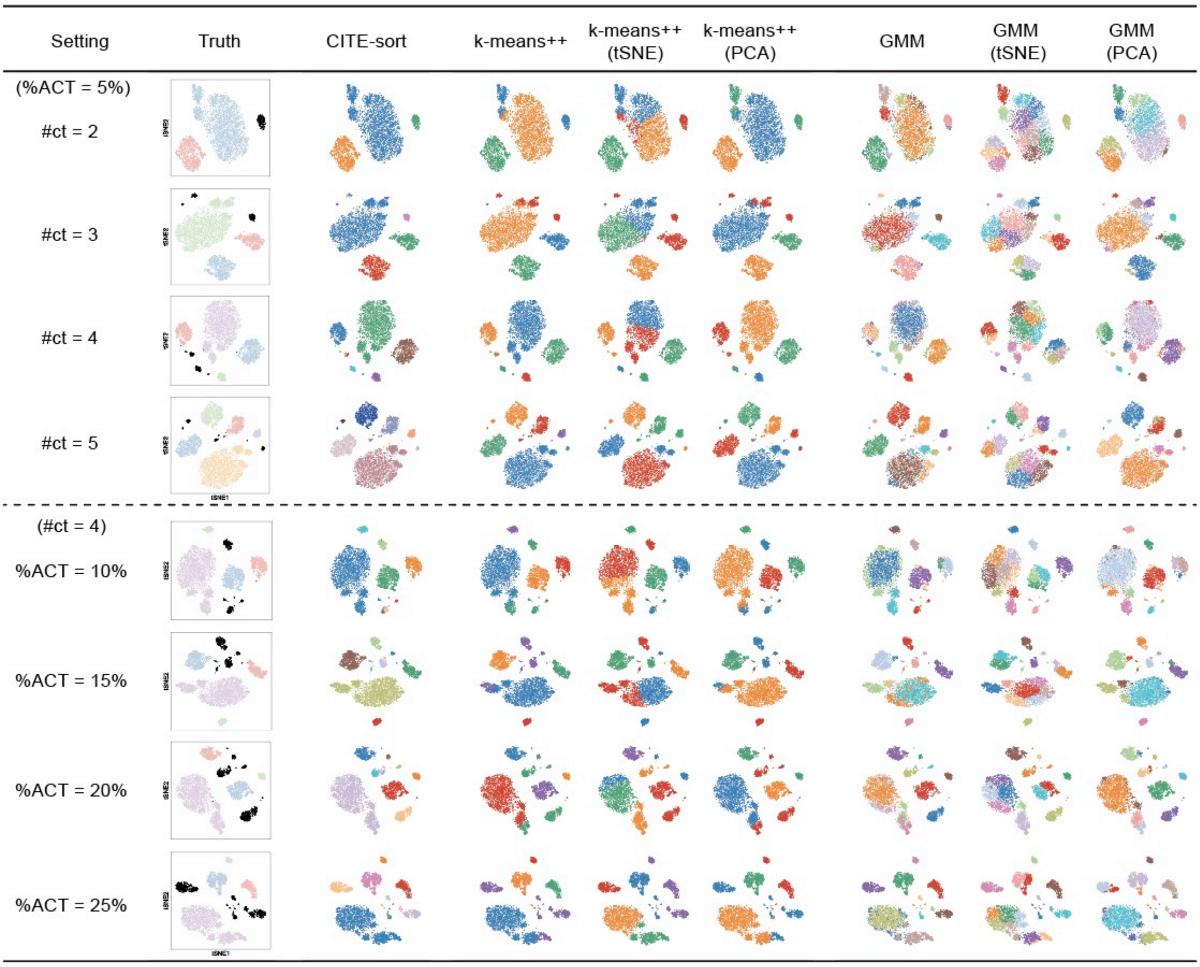


**Supplementary Figure 2**

**Clustering results of CITE-sort and 6 other clustering methods with regard to 8 simulation datasets.**

**Supplementary Table 1**

**S.Table 1.** Hand-curated biological cell types for simulation

| Cell type | Droplet counts |
| --- | --- |
| $CD14^+CD16^-$ | 1972 |
| $CD19^+$ | 418 |
| $CD3^+CD4^+$ | 5803 |
| $CD3^+CD8^+$ | 1965 |
| $CD56^+CD16^+$ | 1290 |

**Supplementary Method 1**

### Benchmarking CITE-sort against GMM and its variants in real datasets

We benchmarked CITE-sort with GMM and its two variants (nGMM and dpgmm) on 8 real CITE-seq datasets. The variant nGMM means that each component has diagonal covariance matrix. GMM and nGMM were implemented by “sklearn.mixture.GaussianMixture” with parameter “covariance_type” set as “full” and “diag”, respectively. The component numbers of GMM and nGMM were determined by Bayesian information criterion (the largest component number was set as 50). The variant dpgmm is variational Bayesian estimation of a Gaussian mixture. It could infer the effective number of components from data. It was implemented by “sklearn.mixture.BayesianGaussianMixture”. For each dataset, we run each method for 10 times and recorded the exculation times and likilihood values. The mean and standard deviation of likelihood and excution time are displayed in Figure 11.
